# Effect of pre-germination temperature conditions on germination characteristics of temperate grassland species

**DOI:** 10.1101/2024.09.26.615133

**Authors:** Réka Kiss, Katalin Lukács, Ágnes Tóth, Benedek Tóth, Kenz Raouf Samraoui, Rita Engel, Balázs Deák, Orsolya Valkó

**Affiliations:** ’Lendület’ Seed Ecology Research Group, Institute of Ecology and Botany, HUN-REN Centre for Ecological Research, Vácrátót, Hungary; National Laboratory for Health Security, HUN-REN Centre for Ecological Research, Budapest, Hungary; University of Szeged, Department of Ecology, Szeged, Hungary; University of South Bohemia, Faculty of Science, České Budějovice, Czechia

**Keywords:** climate change, dormancy, germination capacity, grassland specialist species, stratification

## Abstract

Germination characteristics of target species used for grassland restoration are often unknown, despite such information would be essential for designing restoration interventions. Filling this knowledge gap and improving our understanding about germination responses of grassland species to varying cold and warm treatments could be especially important in the face of climate change. Here we studied the effect of three different durations of warm dry stratification and three different durations of cold wet stratification, and their combinations in a full factorial design (in total 15 different pre-germination treatments), on the germination capacity of 48 grassland species native to Central Europe. Stratification treatments modelled present and forecasted summer (1-3 months warm period) and winter (1-3 months cold period) temperature conditions, while the study of the combined effect of these treatments is especially important in spring-germinating species. As response variables, we calculated relative response indexes and germination uncertainties of each species separately and applied general linear models to study the effect of treatments on these variables. We found clear effect of warm- or cold stratification on relative response indexes only in four species: strong positive response to warm stratification was found in *Silene conica*, while strong positive response to cold stratification was found in *Agrimonia eupatoria, Echium vulgare*, and *Plantago lanceolata*. The responses to treatment combinations were contradictory or lacked clear trends in most of the species. Germination uncertainty in general was high for all species, supporting the fact that Central European grassland species often rely on bet hedging as risk spreading strategy, to avoid unfavourable conditions during seedling establishment.

## Introduction

Grasslands are species-rich communities that are among the most diverse ecosystems in the world (Dengler et al. 2014). Considering their importance in providing various ecosystem services, such as mitigation of the effects of climate change, regulation of water supplies, biomass production, and pollination, their conservation is of great importance (Dass et al. 2018, Dudley et al. 2020, Bardgett et al. 2021). Conservation of grasslands implies the protection of existing ones and the restoration of degraded and vanished ones (Dudley et al. 2020, Tölgyesi et al. 2022). Grassland restoration often requires active interventions including species (re)introduction to the target areas (Hall et al. 2021). Seed sowing is a widely used method for seed introduction; seed mixtures composed of native plants of local provenance proved to efficiently sustain grassland restoration (Shaw et al. 2020, Shackelford et al. 2021, Kaulfuß et al 2022, Kiss et al. 2022). However, the amount of available seeds is limited while the demand for restoration increases continuously (Bardgett et al. 2021, Ladouceur et al. 2018, Török et al. 2024). Therefore, the efficiency of restoration by seed sowing should be increased. This can be either achieved by seed enhancement techniques (i.e. seed priming and coating; Pedrini et al. 2020), or by the conscious seed use (i.e. sowing in adequate period or applying pre-treatments that alleviate dormancy; Kildisheva et al. 2020). A deeper understanding of seed dormancy and germination characteristics of the target species could further improve seed use efficiency. However, despite these characteristics should be considered during restoration projects, the existing knowledge gaps delay their consideration.

Seed germination is induced by a set of environmental cues related to temperature, water availability and light conditions (Baskin & Baskin 2014); upon meeting the adequate conditions seed germination occurs. Favourable conditions often occur in a different period of the year than seed dispersal; and plants overcome the unfavourable conditions of this intermediate period by producing dormant seeds. Seed dormancy is the mechanism that prevents seed germination under unfavourable conditions but also delays germination under apparently favourable conditions (Baskin & Baskin 2004, Jayasuriya et al 2009). The presence of a fraction of seeds which remains dormant under favourable conditions is a risk-spreading strategy of plants; it reduces year-to-year variation in fitness by prolonged seed dormancy and variation in germination timing, a strategy known as bet-hedging (Cohen 1966, Seger & Brockmann 1987, Venable 2007, Ooi et al. 2009, Gremer et al. 2012, Pausas et al. 2022). Dormancy is present in 50–90% of the global wild flora (Baskin & Baskin 2014). The type of dormancy as well as the dormancy breaking mechanism depends on various factors, such as phylogeny, geography, climate, environmental predictability, and habitat type (Cohen 1966, Nikolaeva 2001, Rubio de Casas et al. 2017, Wyse & Dickie 2018, Gioria et al. 2020, Carta et al. 2022, Zhang et al. 2022, Pausas et al. 2022, Rosbakh et al. 2023). There are five major dormancy types and several subtypes; the most common dormancy types are physiological dormancy (PD), i.e., when the embryo possesses an inhibiting mechanism that prevents emergence, and physical dormancy (PY), i.e., when the seed possesses a hard seed coat impenetrable to water (Finch-Savage & Leubner-Metzger 2006, Baskin & Baskin 2004, 2014). Dormancy breaking mechanisms differ between dormancy types: a dry period after seed dispersal is enough for some species (dry after-ripening), while others require stratification (exposed to warm or cold temperature, which are signals of summer or winter periods) or scarification (mechanical injury of the seed coat to allow water uptake) or the combination of different conditions and treatments. The optimal length and strength of these conditions is species specific and if not met, instead of germination they can induce secondary dormancy or loss of seed viability (Reed et al. 2022, Maleki et al. 2023).

Despite their importance, dormancy type and dormancy breaking mechanisms are unknown for many species and many regions (Fernández-Pascual et al. 2023). Besides, intraspecific differences between populations and even individuals also exist, which further deepens the knowledge gap (Baskin & Baskin 2023). This knowledge gap not only decreases restoration efficiency by seed sowing, but it also creates a problem in the face of climate change even in those regions, from where there is more data on the dormancy types of plant species, such as in temperate Europe (Baskin & Baskin 2014, Rosbakh et al. 2020). Climate change is predicted to raise temperatures in Central Europe: the warming will be the greatest in spring periods and lowest in autumns, while great variability in winter warming makes this season very unpredictable (Twardosz et al. 2021). Aside from warming, droughts will become more frequent and more severe especially during growing seasons (Hari et al. 2020, Jaagus et al. 2020). Without the basic knowledge about dormancy and germination characteristics, it is hard to forecast species- and also community-level responses to changing environment (i.e. climate change) and to improve the efficiency of seed-based restoration actions (Walck et al. 2011, Larson et al. 2015, Kövendi-Jakó et al. 2017, Kildisheva et al. 2020).

In our work we aimed to contribute to filling the knowledge gap regarding the dormancy and germination characteristics of Central European plants that are target species in ecological restoration. We selected 48 plant taxa characteristic of various grassland habitats for our study, which enhances our ability to form community-level predictions and design restoration plans in the future. Our aim was to explore the germination response of species to three different duration of warm dry and three different duration of cold wet stratification treatments, and their combinations in a full factorial design (in total 15 different pre-germination treatments). Our questions were: i) What is the effect of the increasing length of the warm period on the germination characteristics of the studied species? ii) What is the effect of the decreasing length of the cold periods on the germination characteristics of the studied species?

## Materials and methods

### Pre-germination treatments

We selected 48 typical grassland species for this experiment, that represent well the target species pool of restoration projects in lowland grasslands in Hungary (Supplementary Table 1). The seeds were purchased from a seed producer of native species who collected and produced the seeds originating in the Kiskunság region (Central Hungary). A batch of 125 seeds of each species (25 seeds in five replicates) was subjected to each stratification treatment (in total 1,875 seeds per species). Stratification treatments were designed as to model the changes in the length of the summer and winter periods: we used one-, two- or three months of warm dry stratification (warm stratification (W), at 24°C), one-, two- or three months of cold wet stratification (cold stratification (C), at 4°C) and the full factorial combination of the treatments (WC). Under natural conditions short-warm periods and longer cold periods were characteristics of the study region in the past; under climate change the length of warm periods increases while that of cold periods decreases (Twardosz et al. 2021, Szabó et al. 2024). The combination of treatments is important especially in case of spring-germinating species, which are affected both by the summer and winter conditions, so changes in both seasons will affect their germination success.

For all pre-germination treatments seeds were placed between two one-layered filter papers in zip-lock bags. In cold stratification treatments filter papers were moistened with distilled water. Zip-lock bags containing the seeds of the 48 species were subjected to complete darkness by wrapping them in two layers aluminium foil. Packs subjected to warm stratification were kept in a dark room while bags subjected to cold stratification were kept in refrigerator during the stratification period and were removed only before the start of the germination experiment.

### Germination experiment

At the end of the stratification periods the five replicates of 25 seeds/species were placed in five Petri dishes (110 mm) in one layer of filter paper moistened with 3-4 ml of distilled water depending on the size of the seeds. Petri dishes were put in growing chamber (PHCbi MLR-352) in 12 h photoperiod on 20/10°C. The germination experiment lasted for four weeks, seedling emergence was verified weekly. Germinated propagules were counted and removed from Petri dishes. As the capacity of the growing chambers was limited, we needed to germinate seeds in four different cycles; the stratification treatments were designed as to their length to match with the germination cycles (Supplementary Table 2). In each germination cycle we also germinated five replicates of 25 untreated seeds (dry storage in room temperature, 20°C) of each species as controls (K).

### Statistical analysis

For the statistical analysis we calculated the Relative Response Index (RRI, Armas et al. 2004) and the germination uncertainty (U, Shannon 1948; Labouriau & Valadares 1976; Labouriau 1983) for each species in treatment separately. RRI shows the effects of the treatments compared to the control and it was calculated as follows:

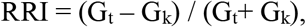

where G_t_ and G_k_ are the number of seedlings germinated in a particular treatment (G_t_) and in the untreated control (G_k_), respectively. RRI ranges between –1 and +1, zero means that the control and the treatment are not different. U measures the degree of germination dispersion, a lower value being associated with a more concentrated germination in time, while a higher number means a prolonged germination period. U was calculated using the *germinationmetrics* package (Aravind et al. 2023); to calculate and compare U, we removed treatments where germination was not detected in given treatments or mean germination was below 4% (i.e. one seed germinated from 25 seeds in a Petri dish, see Supplementary Table 4 and 5). We removed outliers from RRI data and added +2 to each RRI score and +1 to each U to be suitable to analyse it with Gamma distributed generalised linear models (GLMs); we used stratification treatments as explanatory variables (fix factor, 15 levels), RRI and U scores as dependent variables. GLMs with binomial distribution were used to compare mean germination success in the control treatments of the four germination cycles. All statistical analyses were performed in R statistical environment (version 4.3.2, R Core Team 2023). Four species (*Galium boreale, G. verum, Salvia austriaca, Schoenus nigricans*) did not germinate adequately to support their inclusion in statistical analysis, they were removed entirely from the dataset.

## Results

Significant differences between stratification treatments were found in the majority of species (41 species), while treatments had no effect on germination success in three species (DP, PAr, TeC) (Figure 1, Table 1, Supplementary Figure 1). Stratification treatments had significant effect on the U of 31 species and had no effect on the U of 13 species (Figure 1, Supplementary Table 3, Supplementary Figure 1).

**Table 1.** Mean relative response index (RRI) of 44 grassland species in the 15 stratification treatments. The darkening hue of cells indicates the increasing mean RRI, different hues indicate significant differences (*p* ≤ 0.05). Species are abbreviated using the first letters of genus and species names, for the full list of abbreviations, please see Supplementary Table 1. Treatment codes are abbreviated as follows: W1-3 – 1-3 months of warm stratification, C1-3: 1-3 months of cold stratification. For further information, see Supplementary Figure 1.

|  | W1 | W2 | W3 | C1 | C2 | C3 | W1C1 | W1C2 | W1C3 | W2C1 | W2C2 | W2C3 | W3C1 | W3C2 | W3C3 |
| --- | --- | --- | --- | --- | --- | --- | --- | --- | --- | --- | --- | --- | --- | --- | --- |
| AM | 0.04 | 0.03 | 0.05 | -0.01 | -0.02 | 0.09 | 0.03 | 0.07 | 0.02 | 0.08 | 0.00 | 0.06 | -0.01 | 0.05 | 0.05 |
| AE | 0.11 | -0.43 | -0.63 | 0.89 | 0.94 | 0.75 | 0.91 | 0.76 | 0.92 | 0.74 | 0.92 | 0.92 | 0.89 | 0.92 | 0.95 |
| AV | 0.03 | 0.02 | 0.05 | -0.06 | -0.08 | -0.66 | -0.02 | -0.47 | -0.94 | -0.12 | -0.22 | -1.00 | 0.00 | -0.13 | -1.00 |
| AC | -0.04 | -0.02 | 0.02 | 0.15 | 0.02 | -0.02 | 0.17 | -0.18 | -0.03 | -0.05 | -0.04 | 0.10 | 0.19 | 0.24 | -0.11 |
| BO | 0.20 | 0.11 | 0.20 | 0.15 | 0.16 | 0.17 | 0.04 | 0.14 | 0.10 | 0.20 | 0.23 | -0.01 | 0.14 | 0.03 | 0.46 |
| BP | -0.10 | -0.07 | 0.02 | -0.10 | 0.04 | 0.08 | -0.05 | 0.06 | -0.02 | -0.03 | 0.05 | -0.26 | 0.05 | 0.09 | -0.02 |
| CJJ | 0.01 | -0.02 | -0.01 | 0.01 | 0.01 | -0.02 | 0.02 | -0.08 | -0.05 | -0.04 | -0.01 | -0.02 | 0.05 | -0.05 | 0.02 |
| CJP | 0.08 | 0.13 | 0.11 | -0.03 | -0.29 | -0.08 | -0.54 | -0.13 | 0.03 | -0.23 | -0.13 | -0.03 | 0.16 | 0.12 | -0.05 |
| CG | -0.01 | 0.06 | 0.41 | 0.02 | -0.09 | 0.33 | -0.03 | 0.12 | -0.46 | 0.43 | 0.58 | 0.11 | 0.59 | 0.30 | 0.08 |
| DG | -0.02 | 0.09 | 0.11 | -0.04 | 0.08 | 0.13 | 0.06 | 0.13 | 0.09 | 0.14 | -0.01 | 0.12 | 0.04 | 0.14 | 0.13 |
| DP | 0.01 | -0.02 | 0.02 | 0.02 | -0.02 | -0.02 | 0.02 | 0.00 | 0.01 | 0.01 | -0.03 | -0.04 | 0.02 | 0.02 | 0.03 |
| EV | -0.05 | -0.19 | 0.13 | 0.28 | 0.38 | 0.29 | 0.32 | 0.21 | 0.21 | 0.34 | 0.20 | 0.12 | 0.29 | 0.18 | 0.28 |
| FR | 0.16 | 0.14 | 0.25 | 0.14 | 0.17 | 0.06 | 0.12 | 0.18 | 0.15 | 0.27 | 0.22 | 0.12 | 0.23 | 0.26 | 0.00 |
| FVg | -0.01 | -0.06 | 0.29 | -0.02 | -0.12 | 0.25 | -0.12 | 0.04 | 0.03 | 0.26 | 0.20 | 0.24 | 0.11 | 0.23 | 0.15 |
| FW | 0.08 | 0.15 | 0.12 | 0.14 | 0.16 | 0.04 | 0.13 | 0.10 | 0.03 | 0.11 | -0.04 | 0.06 | 0.02 | 0.11 | 0.05 |
| FiV | -0.40 | 0.05 | -0.45 | -0.67 | -0.60 | -0.89 | -0.60 | -0.89 | -1.00 | -0.43 | -0.57 | -1.00 | -0.09 | -0.09 | -1.00 |
| GP | -0.02 | 0.03 | 0.04 | -0.11 | -0.22 | -0.20 | -0.20 | -0.10 | -0.32 | -0.10 | 0.19 | -0.12 | 0.03 | -0.14 | -0.36 |
| HoR | -0.25 | 0.06 | 0.04 | 0.21 | 0.24 | 0.21 | 0.15 | 0.20 | 0.09 | 0.20 | 0.10 | 0.10 | 0.08 | 0.09 | 0.15 |
| HP | -0.07 | 0.07 | 0.00 | 0.04 | -0.02 | -0.10 | 0.02 | 0.03 | 0.02 | 0.03 | 0.04 | -0.16 | -0.29 | 0.02 | -0.07 |
| HM | -0.09 | 0.03 | 0.05 | -0.02 | -0.03 | -0.07 | 0.01 | -0.01 | -0.01 | 0.00 | -0.03 | -0.36 | -0.01 | -0.04 | 0.07 |
| KA | -0.33 | 0.03 | 0.15 | -0.38 | -0.40 | -0.33 | -0.75 | -1.00 | -1.00 | -0.47 | -1.00 | -0.64 | -0.53 | -0.39 | -1.00 |
| LVu | 0.01 | -0.45 | 0.08 | 0.16 | -0.56 | -0.55 | -0.16 | 0.26 | 0.05 | 0.06 | -0.24 | -0.03 | 0.35 | 0.07 | 0.18 |
| LA | 0.04 | 0.03 | 0.05 | -0.15 | -0.03 | -0.09 | -0.07 | -0.05 | -0.20 | -0.06 | -0.06 | -0.10 | -0.09 | -0.04 | -0.20 |
| LC | -0.06 | -0.21 | 0.00 | -0.67 | -0.46 | -1.00 | -0.22 | -0.44 | -0.25 | -0.34 | -0.15 | -0.26 | -0.33 | -0.24 | 0.30 |
| LyV | 0.03 | 0.10 | -0.08 | 0.14 | 0.19 | 0.10 | 0.16 | 0.08 | 0.08 | 0.11 | 0.10 | 0.15 | 0.08 | 0.10 | 0.21 |
| ML | 0.01 | 0.03 | 0.01 | -0.03 | -0.07 | 0.06 | 0.03 | 0.06 | -0.41 | 0.06 | -0.03 | -0.11 | 0.09 | -0.13 | -0.43 |
| OA | -0.02 | -0.04 | 0.05 | 0.01 | -0.06 | -0.16 | -0.05 | -0.11 | -0.33 | 0.03 | -0.06 | -0.19 | -0.02 | -0.05 | -0.58 |
| PL | 0.12 | 0.04 | 0.15 | 0.65 | 0.75 | 0.81 | 0.68 | 0.79 | 0.75 | 0.76 | 0.73 | 0.74 | 0.69 | 0.71 | 0.63 |
| PoA | -0.02 | -0.05 | 0.04 | 0.00 | 0.00 | 0.02 | 0.00 | 0.03 | 0.01 | 0.04 | -0.01 | 0.02 | 0.02 | 0.00 | 0.04 |
| PAr | -0.03 | 0.01 | 0.00 | -0.08 | -0.17 | -0.14 | -0.06 | -0.11 | -0.17 | 0.05 | -0.16 | -0.19 | -0.05 | -0.03 | -0.07 |
| PR | 0.03 | 0.12 | 0.09 | 0.01 | 0.12 | 0.01 | 0.15 | -0.02 | -0.05 | -0.02 | -0.01 | 0.10 | -0.01 | 0.07 | 0.09 |
| SaP | -1.00 | -0.26 | 1.00 | -0.60 | -0.39 | 0.00 | -0.39 | 0.40 | -1.00 | 0.60 | -1.00 | 0.39 | 0.25 | 0.25 | 0.43 |
| SO | -0.02 | -0.11 | 0.29 | -0.02 | -0.29 | 0.41 | -0.27 | 0.10 | 0.42 | 0.17 | 0.23 | -0.28 | 0.34 | 0.34 | -0.55 |
| SJ | 0.11 | -0.14 | 0.00 | 0.04 | -0.12 | -0.01 | -0.01 | -0.28 | 0.20 | -0.16 | 0.02 | 0.03 | 0.21 | 0.38 | 0.16 |
| SC | 0.01 | 0.05 | 0.02 | -0.75 | -1.00 | -1.00 | -0.97 | -1.00 | -1.00 | -1.00 | -1.00 | -1.00 | -0.74 | -1.00 | -1.00 |
| SV | -0.01 | 0.03 | 0.03 | -0.01 | -0.02 | 0.02 | 0.03 | 0.00 | 0.09 | 0.00 | 0.10 | 0.06 | 0.05 | 0.08 | 0.17 |
| StG | -0.46 | -0.29 | -0.60 | 0.26 | -0.10 | 0.08 | -0.25 | 0.01 | -0.22 | -1.00 | 0.02 | -0.17 | -0.18 | -0.17 | 0.31 |
| StP | 0.02 | -0.31 | 0.00 | -0.50 | -0.05 | 0.36 | -0.23 | 0.25 | 0.92 | 0.14 | 0.91 | 0.12 | 0.78 | 0.94 | 1.00 |
| TG | -0.08 | -0.06 | 0.11 | -0.40 | -0.13 | -0.34 | -0.20 | 0.04 | 0.19 | 0.08 | 0.22 | 0.10 | 0.05 | 0.05 | 0.17 |
| TeC | 0.01 | -0.04 | -0.31 | 0.01 | -0.14 | -0.17 | -0.26 | -0.33 | -0.10 | -0.27 | -0.22 | -0.76 | -0.61 | -0.23 | -0.69 |
| TD | 0.06 | -0.09 | 0.05 | -0.19 | -0.18 | -0.01 | -0.01 | -0.11 | 0.52 | -0.03 | 0.44 | 0.46 | 0.44 | 0.50 | 0.37 |
| TM | 0.03 | 0.03 | 0.20 | -0.10 | 0.02 | -0.27 | 0.02 | -0.11 | -0.11 | 0.03 | -0.20 | -0.30 | 0.19 | 0.00 | 0.15 |
| TR | -0.08 | -0.15 | 0.00 | -0.08 | -0.11 | -0.29 | -0.02 | -0.04 | -0.03 | -0.02 | -0.06 | 0.04 | -0.07 | 0.09 | 0.01 |
| VP | -0.02 | -0.02 | 0.02 | -0.02 | -0.06 | 0.02 | -0.01 | 0.00 | -0.01 | -0.01 | -0.03 | -0.03 | -0.01 | -0.02 | -0.02 |

**Figure 1.**
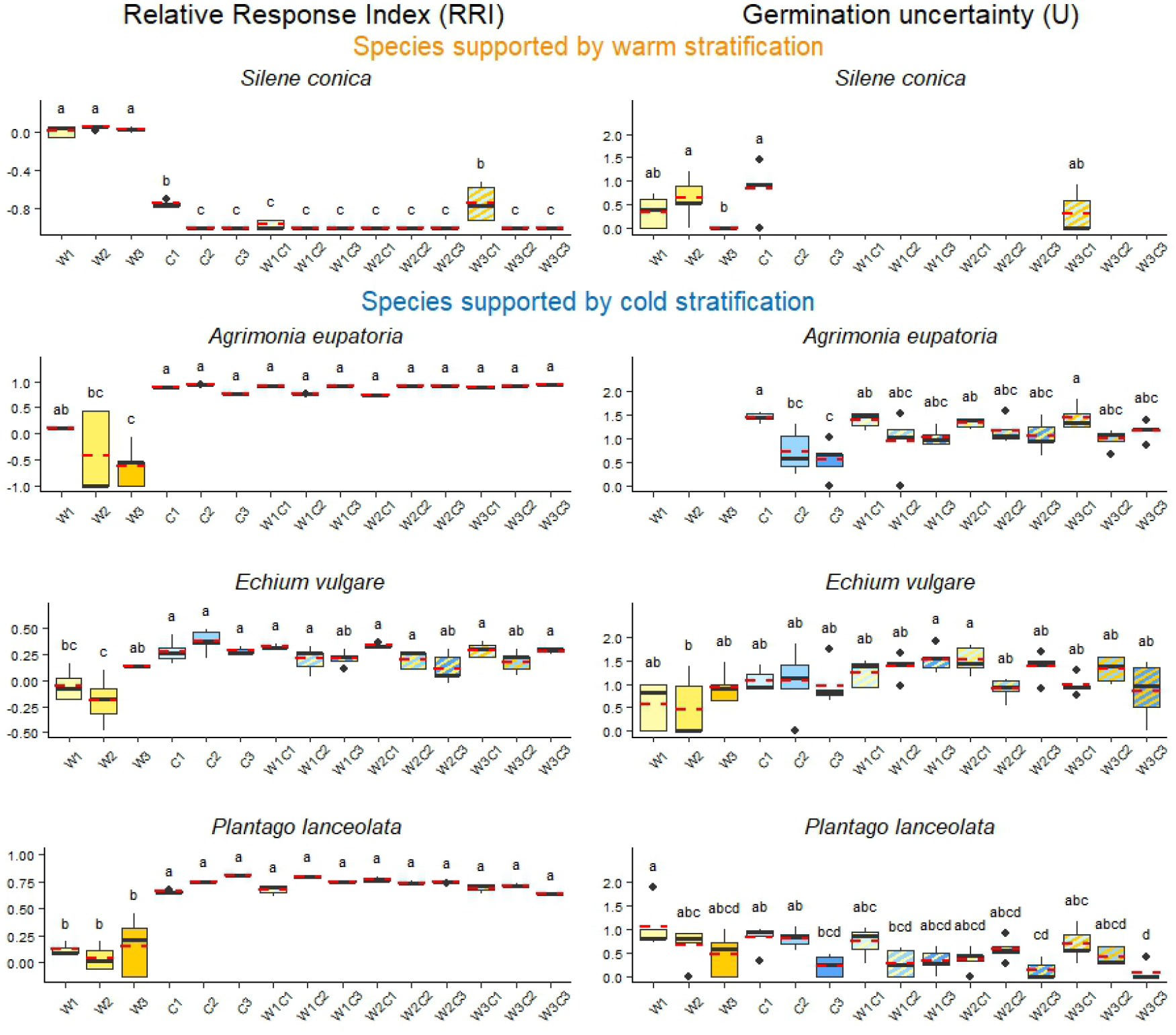
Response in the Relative response index (RRI, y axis of the left column) and Germination uncertainty (U, y axis of the right column) of four model species with clear responses to the stratification treatments. Medians are displayed with solid black lines and means with dashed red lines. Lower-case letters indicate significant differences between the treatments (GLM, p|<0.05). Treatment codes are abbreviated as follows: W1-3: 1-3 months of warm stratification, C1-3: 1-3 months of cold stratification. For the germination responses of all the studied species, see Table 1, Supplementary Figure 1.

Germination success of the controls of the four germination cycles was significantly different in 21 species and was similar in 23 species (Supplementary Table 4, Supplementary Figure 2). Germination uncertainty (U) of the controls of the four germination cycles was significantly different in 8 species and was similar in all other species (36 species) (Supplementary Table 5, Supplementary Figure 2).

### Light requirements of species for germination

There was a proportion of species whose seeds started to germinate already in the dark in the cold wet stratification treatments. We noted these germination events as well, since these gave useful qualitative information on the light requirements of the species during germination. The 16 species which were able to germinate in dark were: *Anthyllis vulneraria, Dactylis glomerata, Dianthus pontederae, Echium vulgare, Festuca rupicola, Festuca vaginata, Festuca wagneri, Gypsophila paniculata, Hypochoeris maculata, Linum austriacum, Lotus corniculatus, Medicago lupulina, Potentilla argentea, Silene vulgaris, Trifolium montanum, Trifolium repens*.

## Discussion

There were only few species which showed clear responses to the warm or cold stratification treatments, but even in these species, the length of these periods was mostly irrelevant. Clear trends in response to warm- or cold stratification were present in RRIs of four species, but it was contradictory or lacked clear trends in the other species. In case of U we found that in general it was high, even when germination success was high, supporting the idea that bet-hedging strategy is generally present in Central European plant species, to minimalize the risks occurring during seedling establishment.

### Effect of the warm periods

A clear, strong positive response to warm stratification was found in *Silene conica*, while in other species there was no clear trend. *Silene conica* expressed high germination success after warm stratification; while one-month cold stratification decreased drastically its germination percentage, and longer cold periods hampered germination totally. Increased germination by warm stratification and decreased one due to cold stratification was found also in other *Silene* species of arid regions with hot, dry summers (Zani & Müller 2018), and it is a characteristic of winter annuals (Baskin & Baskin 1976, Hicks et al. 2019). *S. conica* can be clearly categorized as an autumn germinating species, favouring germination at the end of warm, hot summers, during the favourable growing conditions of the autumn period with increased precipitation level, lower temperature, as well as weaker interspecific competition (Kahmen & Poschlod 2008). Besides, despite the risk of frost damage, seedlings overwintering in rosette form will have a head-start at the end of winters, and can continue their life cycle as soon as temperature starts to increase (Baskin & Baskin 2014, Saarinen et al. 2011).

In two *Festuca* species the effects of warm stratification treatments were reduced by the presence of cold period in specific cases, when warm stratification lasted only one month (*Festuca wagneri*) or three months (*F. rupicola*). This result is in contrast with the findings of Kövendi-Jakó et al. (2017): according to their results, one-month long cold stratification under similar conditions already decreased germination success of *F. rupicola and F. vaginata*. However, in our study only longer cold periods had a negative effect on germination success, or, in case of *F. vaginata*, the effect was even positive. Besides, it seems that in *F. vaginata* the effects of warm- and cold stratifications are additive, or secondary dormancy does not occur and the seeds of the species tend to germinate especially after receiving at least a two-month-long warm stratification.

For two species with a wide distribution range (*Knautia arvensis, Linaria vulgaris*) we found that the effect of stratification treatments on germination differed from that found in other studies. We found that warm stratification supported germination of *K. arvensis* more than cold stratification; in contrast, in northern populations of *K. arvensis* a two-month long cold stratification was the one that enhanced the seed germination (Vandvik & Vange 2003). Similarly, *Linaria vulgaris* also showed differences in its germination characteristics compared to the seeds originating from coastal dune habitats of the Baltic region (Necajeva & Probert 2011). In our case germination success decreased with the length of the cold period, while in the Baltic region highest germination of *L. vulgaris* was acquired after 5 months of cold stratification. These results highlight the role of germination characteristics in species distribution: heterogeneous seeds with wide germination niches and variable germination characteristics support the wide distribution of species (Vandvik & Vange 2003, Necajeva & Probert 2011).

### Effect of the cold periods/

Strong positive response to cold stratification, independently of its length, was found in three species: *Agrimonia eupatoria, Echium vulgare*, and *Plantago lanceolata*. The germination success of these species was always higher when cold stratification was applied than in warm stratification only treatments. Warm stratification was either ineffective to break seed-dormancy of these species or induced secondary dormancy. Our results are in agreement with other studies on *A. eupatoria* and *P. lanceolata*, which also found that the seeds of these species germinated better following exposure to cold periods or lower germination temperatures (Mira et al. 2018, Asaadi 2022). Dormancy breaks successfully only after the seeds receive the optimal temperature conditions, i.e., optimal temperature range and optimal duration of this temperature. The optimal temperature levels and duration are species-specific and ranges from few days to few months (Walck et al. 2011, Baskin & Baskin 2014, Krock et al. 2016). In our case a one-month-long cold period was enough for all three species to break dormancy. For *A. eupatoria* shorter periods were also reported, and it can be assumed that the optimal duration of cold temperature needed by this species is between two days (Saffari et al. 2020) and ten days (Asaadi 2022). As the need for cold period is so short, germination can be induced during mild periods or heat waves following cold periods during winter. Seedlings in these periods may be exposed to the risk of frost damage, but overwintering individuals can benefit from favourable growing conditions even shortly after germination. However, to diminish the risks of winter frost periods, seed germination still remains scattered in time, assuring a fraction of seeds to be available for germination later during the growing period.

Germination follows a similar trend to the one described above in *Holoschoenus romanus* and *Lythrum virgatum* as well. In *L. virgatum* not only germination success, but germination synchrony as well was increased by the presence of a cold treatment. The strategy of this species differs from the majority of species; *L. virgatum* focuses on germination during favourable conditions rather than on risk-spreading by bet-hedging. However, considering that *L. virgatum* produces a large quantity of seeds, the persistence of the population can rely on few successfully surviving seeds and seedlings. Contrary to the aforementioned species, where the length of cold period was not important, the length of the cold periods was important in *Stachys palustris* and *Silene vulgaris*, especially when preceded by warm stratification. In both cases it was already known, that cold stratification increases germination success (Kövendi-Jakó et al. 2017), but we also observed that in *S. palustris* the warm and cold stratification treatments expressed an additive effect on germination.

In several species decreased germination was observed in response to prolonged cold periods (*Anthyllis vulneraria, Trifolium repens*) or in treatment combinations where prolonged cold periods followed warm periods (*Filipendula vulgaris, Medicago lupulina, Onobrychis arenaria*). The result of *A. vulneraria* is in accordance with a previous study, which found that the species reacts well to shorter periods with cold temperature (1,5 month in 5ºC, Szalai & Ferschl 2016). In *F. vulgaris* it was demonstrated, that the species germinates better after warmer conditions during seed dormancy (Wagner et al. 2011). In four of the species listed above we can assume that the responses found are caused by complex mechanisms. These species belong to *Fabaceae* family, which often possess physical dormancy or combinational dormancy (Baskin & Baskin 2004): due to their hard, water-impermeable seed coat they are physically dormant, and in some genera the development of embryo is also physiologically inhibited. Germination of the seeds of these species depends both on their ability to uptake water and on the dormancy breaking of the embryo. Sensitivity of the embryo for breaking dormancy is cyclic (Van Assche et al. 2003, Jayasuriya et al. 2009): if not in a sensitive phase, despite water uptake takes place, seeds swell but germination does not occur. The mechanism works in the other way too: seeds may be in a sensitive state to break dormancy, but if water uptake is not possible, germination is hampered and seeds become insensitive again (Jayasuriya et al. 2009). The role of this complex dormancy-breaking mechanism is the same as that of dormancy cycling in species with physiological dormancy: initiates germination only when conditions are favourable for germination, seedling establishment, and survival (Jayasuriya et al 2009).

The length of cold periods, as well that of warm periods, were also important in *Tragopogon dubius*. The effects of the warm and cold stratification treatments were combined in an interesting way: shorter warm period needed to be combined with a longer cold period for successful germination, and with the increasing length of warm period the length of cold period needed for successful germination decreases. This dormancy breaking mechanism was described for seeds possessing intermediate regular physiological dormancy (PD) but it was not known to be present in any of the genera of the *Asteraceae* family (Baskin & Baskin 2023).

#### Unaffected germination patterns

*Dianthus pontederae, Potentilla argentea* and *Teucrium chamaedrys* did not have significant differences in their RRIs in response to the stratification treatments. *D. pontederae* and *P. argentea* had a high germination success even in their controls, which means that either their dormancy was broken during the dry storage, or they have non-deep dormancy and can germinate under various settings as soon as the conditions become favourable. Besides, none of the stratification treatments induced secondary dormancy of the seeds. In a previous study we found similarly high germination success in *D. pontederae* (Kiss et al. 2018), which supports the fact of *D. pontederae* seeds possessing non-deep dormancy and can germinate fast in response to favourable environmental conditions. In contrast *T. chamaedrys* germinated poorly even in controls. Considering that higher germination success for *T. chamaedrys* was reported earlier (Dönmez & Önal 2023), we suspect that our results were caused by the low seed viability in the used seed stock. This explanation can also be applied to *Galium verum*, one of the four species excluded from analysis due to the lack of germination, as our previous study reported high germination capacity of *G. verum* (Kiss et al. 2018). Differences may be resulted from the inter-annual variability of environmental conditions, which influence growing conditions and fitness of mother plants (Giménez-Benavides et al. 2005, Jaganathan 2016). Besides, differences in germination within species can also be found between populations and even within-individual differences were described (Fenner 1991, Mondoni et al. 2014, Penfield & MacGregor 2017, Baskin & Baskin 2023).

### Limitations of the study and further possibilities

Due to the high number of species and treatments and limited space availability in the growing chambers, germination of seeds took place in four consecutive cycles. For each germination event a separate control was included. When we compared the germination success of dry-stored seeds, we found that in half of the species germination success decreased with the length of storage time, possibly as a result of viability-loss or of a prolonged after-ripening period causing secondary dormancy (Walck et al. 2011, Baskin & Baskin 2020, Maleki et al. 2023). To account for the potential decreased germination of seeds, we used the relative response index (RRI), which takes into account the results of controls to calculate response to treatments. We are also aware, that in the present study only the effect of the temperature of the pre-germination period was studied on germination characteristics, while moisture content and temperature during germination was constant. In the future different environmental cues (moisture content, temperature during germination) should be studied both under controlled (laboratory) and under field conditions. Here we only studied germination capacity, but in future studies it would be important to evaluate the fitness of the germinated seedlings.

## Conclusions and implication for restoration

Understanding environmental conditions that induce or hamper germination by inducing or releasing dormancy of wild grassland species has practical implications. Seed-sowing based restoration practices should take into account the species-specific seed dormancy type and use appropriate sowing time combined with dormancy-breaking pre-treatments to increase restoration efficiency. This is of great importance especially in the face of the climate change, when well known environmental conditions change, and therefore restoration measures should be flexibly adapted to the changing climate. We found that in general the grassland species included in our study can cope with climate warming in the phenological stage of germination. Although their aboveground abundance will show great year-to-year variability, their seeds will be able to germinate thanks to the bet-hedging strategy which can support the persistence of the populations.

## Acknowledgements

We are grateful for Eszter Korom, who provided considerable help during the preparation and execution of the experiment. We are thankful to Bernadett Kolonics and Richárdné Ribai who helped in smoothly running of the germination cycles. The authors were supported by the Hungarian National Research, Development and Innovation Office (Grant Numbers: PD 137632 – R.K., KKP 144096 – O.V., FK 135329 – B.D.).

