## Supplementary Material for "Effect of pre-germination temperature conditions on germination characteristics of temperate grassland species"

Supplementary Table 1. List of the studied species. Strikethrough letters indicate species which were excluded from statistical analysis due to the lack of germination.

| Species name | Abbreviation |  | Species name | Abbreviation |
| --- | --- | --- | --- | --- |
| *Achillea millefolium* | AM |  | *Linum austriacum* | LA |
| *Agrimonia eupatoria* | AE |  | *Lotus corniculatus* | LC |
| *Anthyllis vulneraria* | AV |  | *Lythrum virgatum* | LyV |
| *Astragalus cicer* | AC |  | *Medicago lupulina* | ML |
| *Betonica officinalis* | BO |  | *Onobrychis arenaria* | OA |
| *Brachypodium pinnatum* | BP |  | *Plantago lanceolata* | PL |
| *Centaurea jacea* ssp. *jacea* | CJJ |  | *Poa angustifolia* | PoA |
| *Centaurea jacea* ssp. *pannonica* | CJP |  | *Potentilla argentea* | PAr |
| *Chrysopogon gryllus* | CG |  | *Potentilla recta* | PR |
| *Dactylis glomerata* | DG |  | *~~Salvia austriaca~~* | ~~SA~~ |
| *Dianthus pontederae* | DP |  | *Salvia pratensis* | SaP |
| *Echium vulgare* | EV |  | *Scabiosa ochroleuca* | SO |
| *Festuca rupicola* | FR |  | *~~Schoenus nigricans~~* | ~~SN~~ |
| *Festuca vaginata* | FVg |  | *Senecio jacobaea* | SJ |
| *Festuca wagneri* | FW |  | *Silene conica* | SC |
| *Filipendula vulgaris* | FiV |  | *Silene vulgaris* | SV |
| *~~Galium boreale~~* | ~~GB~~ |  | *Stachys germanica* | StG |
| *~~Galium verum~~* | ~~GV~~ |  | *Stachys palustris* | StP |
| *Gypsophila paniculata* | GP |  | *Teucrium chamaedrys* | TeC |
| *Holoschoenus romanus* | HoR |  | *Thymus glabrescens* | TG |
| *Hypericum perforatum* | HP |  | *Tragopogon dubius* | TD |
| *Hypochoeris maculata* | HM |  | *Trifolium montanum* | TM |
| *Knautia arvensis* | KA |  | *Trifolium repens* | TR |
| *Linaria vulgaris* | LVu |  | *Verbascum phoeniceum* | VP |

Supplementary Table 2. Design of the experiment. Orange colour indicates warm stratification, blue colour indicates cold stratification, green colour indicates the germination experiment. Abbreviations: Mo.=Month. Treatment codes are abbreviated as follows: W1-3 – 1-3 months of warm stratification, C1-3: 1-3 months of cold stratification, K – control.

| Code | Stratification | Mo. 1 | Mo. 2 | Mo. 3 | Mo. 4 | Mo. 5 | Mo. 6 | Mo. 7 |
| --- | --- | --- | --- | --- | --- | --- | --- | --- |
| K1 | 0 Mo.,  Control of cycle I |  |  |  |  |  |  |  |
| W1 | 1 Mo. warm |  |  |  |  |  |  |  |
| W2 | 2 Mo. warm |  |  |  |  |  |  |  |
| C1 | 1 Mo. cold |  |  |  |  |  |  |  |
| C2 | 2 Mo. cold |  |  |  |  |  |  |  |
| W1C1 | 1 Mo. warm +  1 Mo. cold |  |  |  |  |  |  |  |
| K2 | 0 Mo. ,  Control of cycle II |  |  |  |  |  |  |  |
| W3 | 3 Mo. warm |  |  |  |  |  |  |  |
| C3 | 3 Mo. cold |  |  |  |  |  |  |  |
| W1C2 | 1 Mo. warm +  2 Mo. cold |  |  |  |  |  |  |  |
| W2C1 | 2 Mo. warm + 1 Mo. cold |  |  |  |  |  |  |  |
| K3 | 0 Mo.,  Control of cycle III |  |  |  |  |  |  |  |
| W1C3 | 1 Mo. warm +  3 Mo. cold |  |  |  |  |  |  |  |
| W2C2 | 2 Mo. warm + 2 Mo. cold |  |  |  |  |  |  |  |
| W2C3 | 2 Mo. warm + 3 Mo. cold |  |  |  |  |  |  |  |
| W3C1 | 3 Mo. warm + 1 Mo. cold |  |  |  |  |  |  |  |
| W3C2 | 3 Mo. warm + 2 Mo. cold |  |  |  |  |  |  |  |
| K4 | 0 Mo.,  Control of cycle IV |  |  |  |  |  |  |  |
| W3C3 | 3 Mo. warm + 3 Mo. cold |  |  |  |  |  |  |  |

Supplementary Table 3. Mean germination uncertainty (U) of the studied species in stratification treatments. The darkening hues of cells indicates the increasing mean U, different hues indicate significant differences (*p* ≤ 0.05). Species are abbreviated using the first letters of genus and species names, for the full list of abbreviations, please see Appendix 1. Treatment codes are abbreviated as follows: W1-3: 1-3 months of warm stratification, C1-3: 1-3 months of cold stratification. NA in the cells indicate the case when germination was not sufficient to calculate U.

|  | W1 | W2 | W3 | C1 | C2 | C3 | W1C1 | W1C2 | W1C3 | W2C1 | W2C2 | W2C3 | W3C1 | W3C2 | W3C3 |
| --- | --- | --- | --- | --- | --- | --- | --- | --- | --- | --- | --- | --- | --- | --- | --- |
| AM | 0.13 | 0.00 | 0.09 | 0.00 | 0.05 | 0.00 | 0.00 | 0.00 | 0.05 | 0.00 | 0.00 | 0.00 | 0.14 | 0.00 | 0.00 |
| AE | NA | NA | NA | 1.46 | 0.72 | 0.55 | 1.40 | 0.95 | 1.03 | 1.34 | 1.16 | 1.06 | 1.44 | 0.99 | 1.17 |
| AV | 0.43 | 0.44 | 0.13 | 0.00 | 0.17 | 0.46 | 0.00 | 0.14 | NA | 0.07 | 0.48 | NA | 0.94 | 0.75 | NA |
| AC | 1.25 | 1.13 | 1.15 | 0.59 | 0.23 | 0.00 | 0.09 | 0.14 | 0.09 | 0.00 | 0.00 | 0.71 | 0.38 | 0.16 | 0.25 |
| BO | 0.94 | 1.09 | 1.49 | 1.27 | 1.02 | 0.85 | 1.30 | 1.10 | 1.29 | 1.18 | 1.04 | 1.30 | 1.00 | 1.03 | 1.21 |
| BP | 1.24 | 1.66 | 1.32 | 1.23 | 0.96 | 0.49 | 1.30 | 0.93 | 0.66 | 1.27 | 1.23 | 1.36 | 0.79 | 1.37 | 1.35 |
| CJJ | 0.91 | 0.93 | 0.60 | 0.54 | 0.21 | 0.51 | 0.56 | 0.63 | 0.78 | 0.55 | 0.34 | 0.63 | 0.38 | 0.67 | 0.41 |
| CJP | 1.00 | 1.13 | 0.45 | 0.58 | 0.51 | 0.58 | 0.33 | 0.16 | 0.34 | 0.53 | 0.50 | 0.33 | 0.78 | 1.00 | 0.16 |
| CG | 1.09 | 1.11 | 0.73 | 1.15 | 0.86 | 0.78 | 0.91 | 0.37 | NA | 0.85 | 0.55 | 0.00 | 1.08 | 0.40 | 0.36 |
| DG | 0.91 | 1.23 | 1.02 | 1.13 | 1.08 | 0.56 | 1.07 | 1.07 | 0.66 | 0.97 | 1.64 | 0.97 | 1.04 | 1.54 | 1.13 |
| DP | 0.28 | 0.14 | 0.00 | 0.69 | 0.33 | 0.84 | 1.00 | 0.29 | 0.91 | 0.80 | 0.95 | 0.85 | 0.82 | 1.00 | 1.05 |
| EV | 0.56 | 0.47 | 0.94 | 1.08 | 1.06 | 0.97 | 1.24 | 1.37 | 1.53 | 1.51 | 0.90 | 1.39 | 0.98 | 1.32 | 0.86 |
| FR | 0.76 | 1.05 | 1.20 | 1.19 | 1.47 | 1.44 | 1.02 | 0.99 | 1.47 | 0.66 | 1.58 | 1.56 | 0.61 | 1.67 | 1.29 |
| FVg | 1.36 | 0.96 | 1.18 | 1.21 | 1.14 | 1.56 | 1.35 | 1.59 | 1.42 | 1.48 | 1.45 | 1.39 | 1.15 | 1.56 | 1.83 |
| FW | 0.97 | 1.02 | 0.78 | 0.98 | 1.27 | 1.87 | 1.28 | 1.28 | 1.28 | 1.26 | 1.77 | 1.70 | 0.94 | 2.07 | 1.59 |
| FiV | 0.34 | 0.71 | 0.35 | NA | NA | NA | NA | NA | NA | 0.73 | NA | NA | NA | NA | NA |
| GP | 0.50 | 0.23 | 0.26 | 0.37 | 0.34 | 0.44 | 0.16 | 0.58 | 0.27 | 0.55 | 0.91 | 1.30 | 0.91 | 0.66 | 0.83 |
| HoR | 0.79 | 1.30 | 1.05 | 1.10 | 0.91 | 1.15 | 1.11 | 0.84 | 0.95 | 0.86 | 0.65 | 0.50 | 0.96 | 0.80 | 0.70 |
| HP | 1.02 | 0.96 | 1.43 | 1.45 | 1.00 | 1.29 | 1.19 | 1.14 | 1.66 | 1.19 | 1.02 | 0.52 | 0.80 | 1.23 | 1.39 |
| HM | 0.80 | 0.88 | 0.99 | 1.19 | 0.72 | 0.29 | 0.88 | 0.18 | 0.40 | 0.11 | 0.27 | 0.90 | 0.56 | 0.84 | 0.37 |
| KA | 0.88 | 0.85 | 0.91 | 0.60 | 0.18 | NA | NA | NA | NA | NA | NA | NA | NA | NA | NA |
| LVu | 0.47 | 0.48 | 0.62 | 0.69 | 0.18 | 0.50 | 0.64 | 0.69 | 1.08 | 0.83 | 0.00 | 0.33 | 1.05 | 0.51 | 0.79 |
| LA | 1.07 | 0.94 | 1.22 | 0.55 | 0.17 | 1.01 | 0.27 | 0.39 | 0.83 | 0.82 | 1.20 | 1.03 | 0.94 | 0.57 | 1.45 |
| LC | 1.21 | 0.82 | 0.25 | 0.00 | 0.00 | 0.00 | 0.76 | 0.71 | 0.26 | 0.00 | 0.60 | 0.18 | 0.16 | 0.26 | 0.06 |
| LyV | 1.28 | 1.26 | 1.14 | 0.00 | 0.37 | 0.05 | 0.20 | 0.06 | 0.00 | 0.13 | 0.46 | 0.00 | 0.09 | 0.83 | 0.05 |
| ML | 0.28 | 0.29 | 0.05 | 1.27 | 1.00 | 0.00 | 0.15 | 0.13 | 0.25 | 0.28 | 1.01 | 1.10 | 0.00 | 0.30 | 0.19 |
| OA | 1.13 | 0.85 | 0.73 | 0.19 | 0.72 | 0.16 | 0.17 | 0.47 | 0.53 | 0.30 | 0.93 | 0.81 | 0.70 | 0.36 | 0.75 |
| PL | 1.05 | 0.67 | 0.46 | 0.83 | 0.81 | 0.23 | 0.74 | 0.28 | 0.33 | 0.37 | 0.58 | 0.14 | 0.69 | 0.43 | 0.09 |
| PoA | 1.03 | 1.08 | 0.96 | 0.93 | 0.95 | 1.62 | 0.77 | 1.04 | 1.58 | 1.16 | 1.37 | 1.41 | 1.08 | 1.31 | 1.27 |
| PAr | 1.01 | 1.00 | 0.58 | 0.88 | 0.96 | 0.31 | 0.94 | 0.50 | 1.08 | 0.55 | 1.07 | 1.38 | 0.94 | 1.17 | 1.08 |
| PR | 1.02 | 0.63 | 0.62 | 0.59 | 0.58 | 0.36 | 0.76 | 0.29 | 0.42 | 0.48 | 0.57 | 0.40 | 0.49 | 0.87 | 0.09 |
| SaP | NA | 0.18 | 0.16 | NA | NA | NA | NA | NA | NA | NA | NA | 0.20 | NA | 0.20 | NA |
| SO | 0.56 | 0.83 | 0.88 | 0.73 | 0.98 | 0.61 | 0.80 | 0.67 | 0.38 | 1.32 | 0.82 | 0.20 | 0.55 | 1.00 | 0.20 |
| SJ | 0.82 | 0.49 | 0.74 | 0.52 | 0.00 | 0.26 | 0.61 | 0.34 | 0.14 | 0.58 | 0.59 | 0.52 | 0.50 | 0.00 | 0.31 |
| SC | 0.35 | 0.63 | 0.00 | 0.84 | NA | NA | NA | NA | NA | NA | NA | NA | 0.30 | NA | NA |
| SV | 0.86 | 0.79 | 0.58 | 0.66 | 0.62 | 0.41 | 0.36 | 0.34 | 0.42 | 0.37 | 0.44 | 0.39 | 0.65 | 0.87 | 0.75 |
| StG | 0.20 | 0.19 | NA | 0.48 | 0.55 | 0.30 | 0.00 | 0.52 | 0.50 | NA | 0.38 | 0.34 | 0.55 | 0.38 | 0.72 |
| StP | 1.05 | 0.82 | 0.93 | 0.62 | 0.79 | 1.32 | 1.12 | 1.39 | 0.83 | 0.89 | 0.78 | 0.68 | 0.50 | 1.04 | 1.01 |
| TeC | 0.40 | 0.64 | 0.58 | 0.52 | 0.30 | 0.65 | 0.55 | 0.32 | 0.54 | 0.83 | 0.77 | NA | NA | 0.93 | NA |
| TG | 1.00 | 0.87 | 0.22 | 0.18 | 0.68 | 0.13 | 0.21 | 0.28 | 0.18 | 0.08 | 0.09 | 0.31 | 0.25 | 0.29 | 0.14 |
| TD | 1.15 | 1.22 | 0.51 | 1.55 | 1.31 | 1.49 | 0.75 | 1.39 | 1.17 | 0.61 | 0.93 | 1.59 | 1.11 | 0.86 | 1.71 |
| TM | 0.59 | 0.47 | 0.07 | 0.99 | 0.14 | 0.00 | 1.06 | 0.00 | 0.00 | 0.94 | 0.28 | 0.34 | 1.07 | 0.65 | 0.42 |
| TR | 0.64 | 0.31 | 0.06 | 1.14 | 0.84 | 0.72 | 1.08 | 0.79 | 0.24 | 0.79 | 0.80 | 0.28 | 1.00 | 0.80 | 0.29 |
| VP | 1.05 | 0.50 | 0.05 | 0.05 | 0.05 | 0.05 | 0.34 | 0.21 | 0.13 | 0.00 | 0.42 | 0.11 | 0.29 | 0.24 | 0.24 |

Supplementary Figure 1. Relative response index (RRI, y axis of first column) and Germination uncertainty (U), y axis of second column) of the species in the 15 treatments. Medians are displayed with solid black lines and means with dashed red lines. Lower-case letters indicate significant differences between the treatments (GLM, p<0.05). Treatment codes are abbreviated as follows: W1-3: 1-3 months of warm stratification, C1-3: 1-3 months of cold stratification.

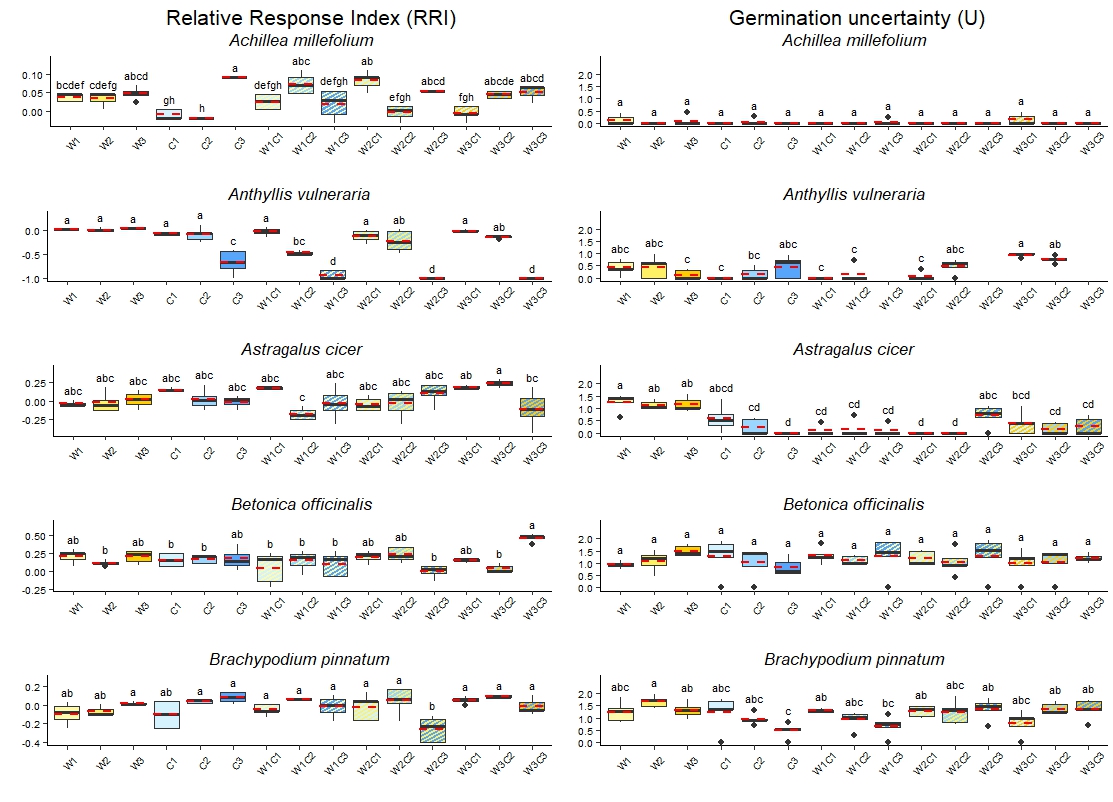

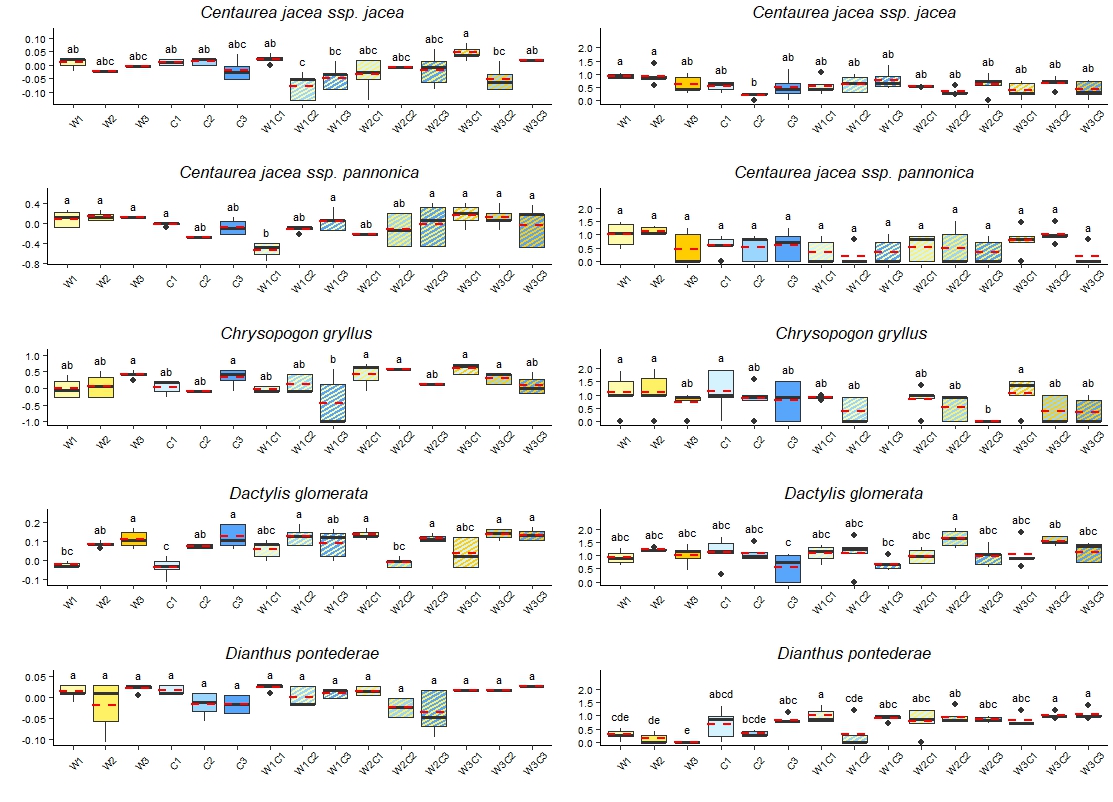

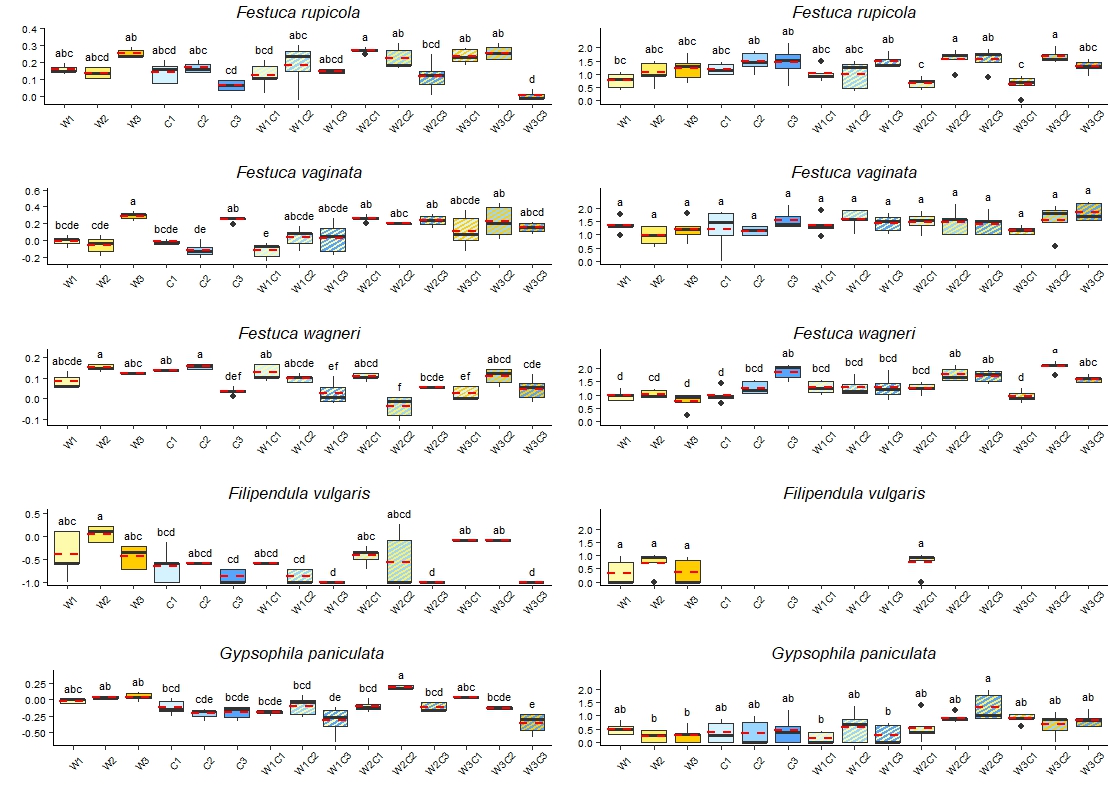

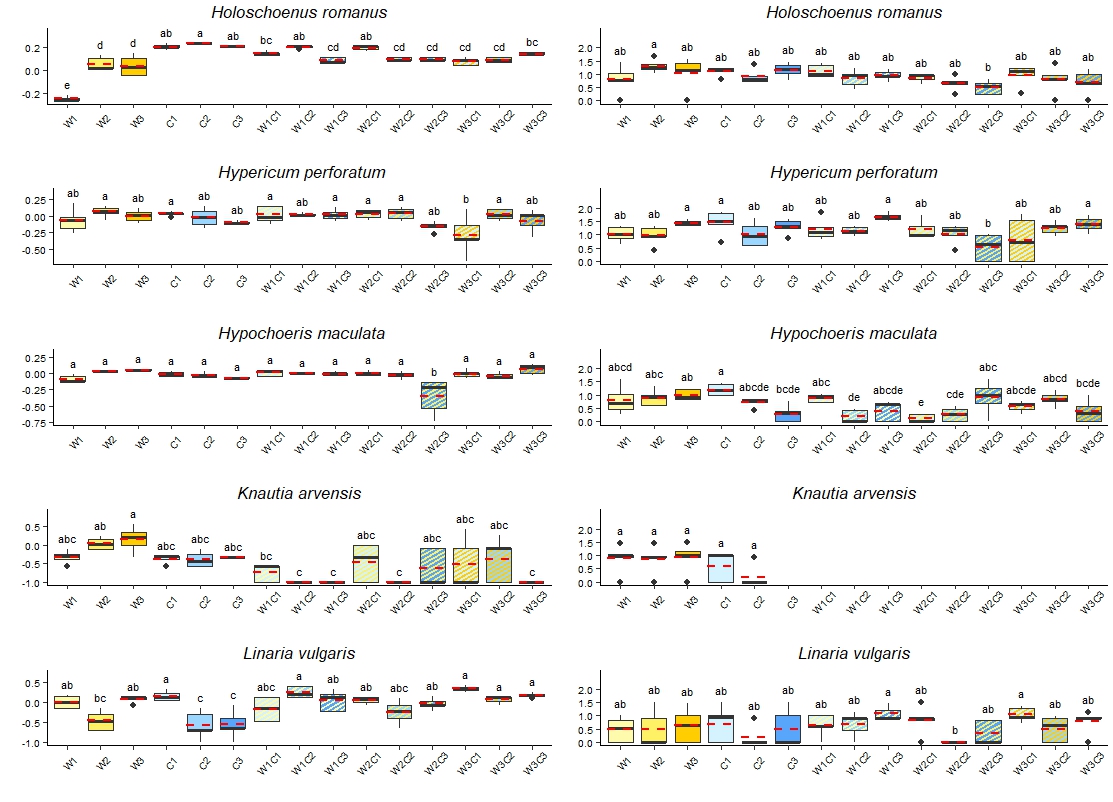

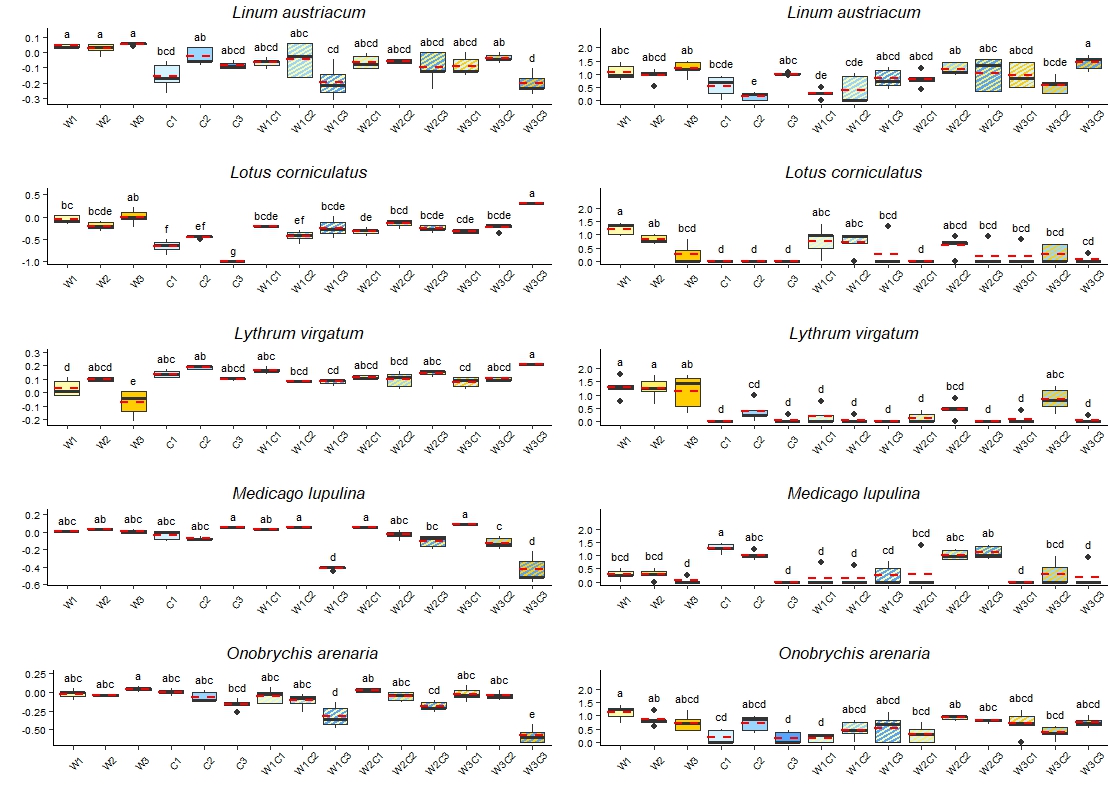

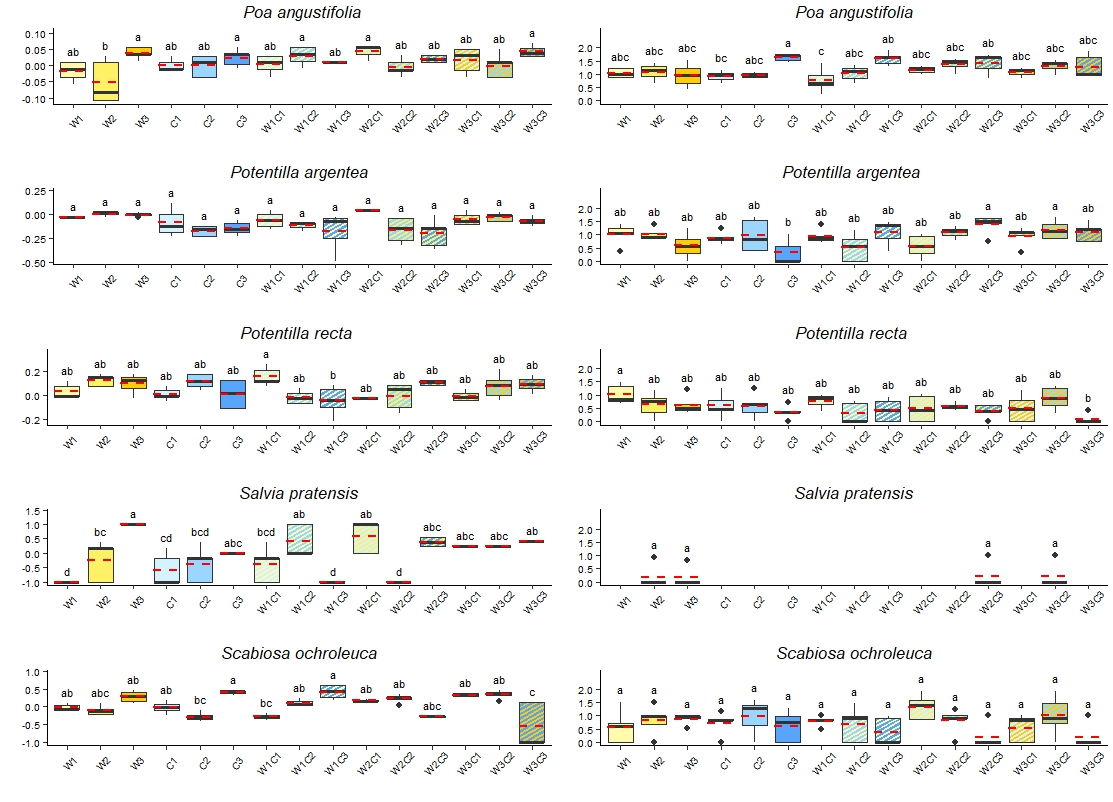

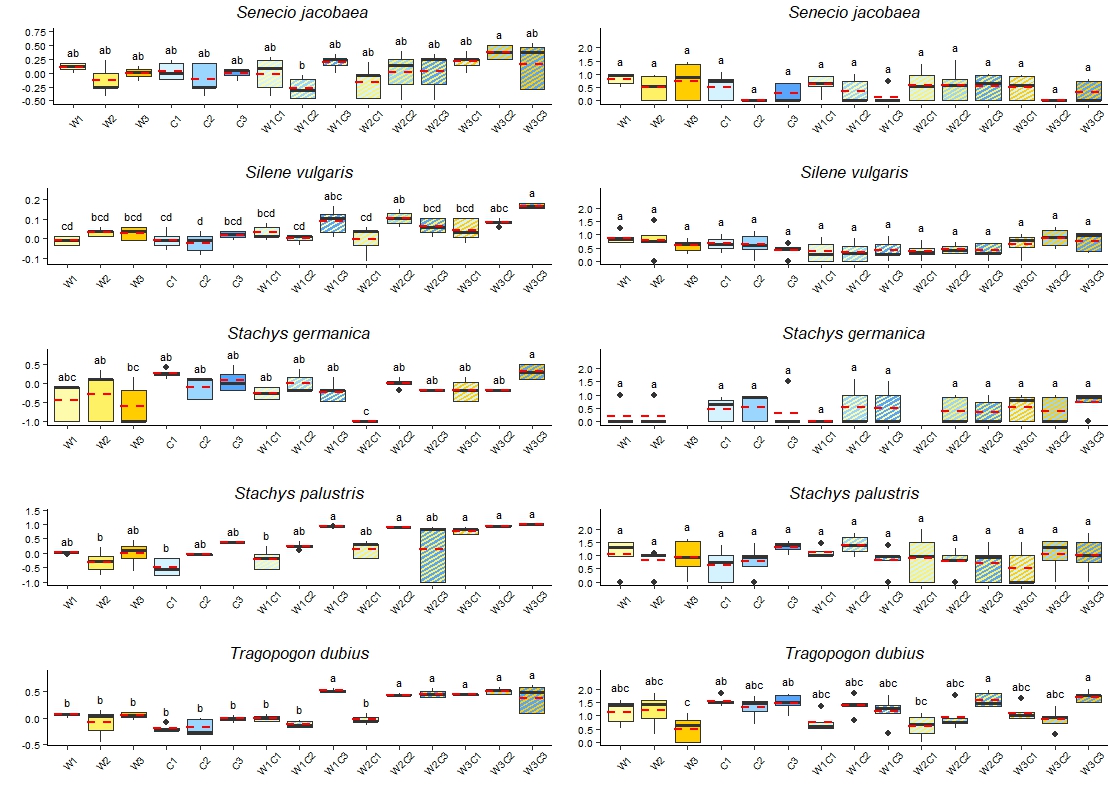

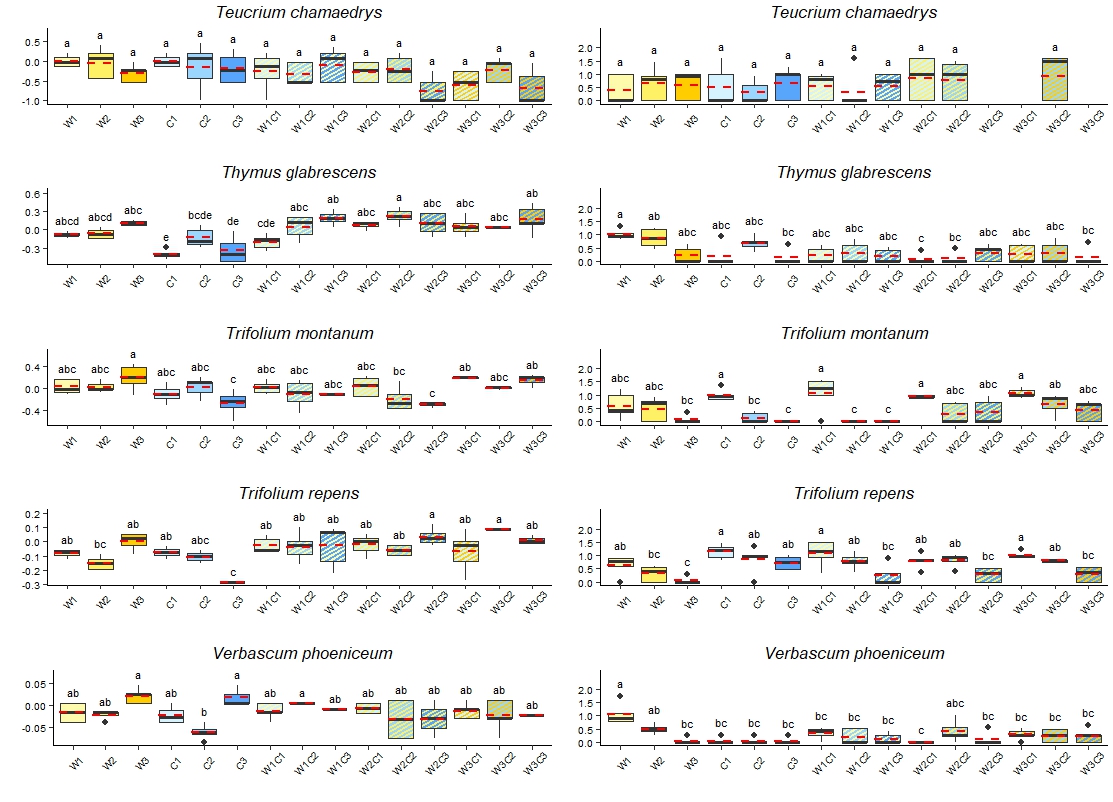

Supplementary Table 4. Mean germination percentage (%) of the controls (K1-K4) of the four germination cycles. The darkening hue of cells indicates the increasing mean germination success, different hues indicate significant differences (*p* ≤ 0.05).

| Species name | Abbreviation | K1 | K2 | K3 | K4 |
| --- | --- | --- | --- | --- | --- |
| *Achillea millefolium* | AM | 91.20 | 80.00 | 89.60 | 88.00 |
| *Agrimonia eupatoria* | AE | 3.20 | 13.60 | 4.00 | 2.40 |
| *Anthyllis vulneraria* | AV | 52.80 | 58.40 | 45.60 | 39.20 |
| *Astragalus cicer* | AC | 35.79 | 41.60 | 30.89 | 30.40 |
| *Betonica officinalis* | BO | 32.00 | 27.20 | 32.00 | 18.40 |
| *Brachypodium pinnatum* | BP | 82.73 | 67.00 | 70.61 | 67.62 |
| *Centaurea jacea* ssp. *jacea* | CJJ | 88.00 | 88.80 | 81.60 | 73.60 |
| *Centaurea jacea* ssp. *pannonica* | CJP | 28.80 | 19.20 | 10.62 | 11.20 |
| *Chrysopogon gryllus* | CG | 14.40 | 4.80 | 3.20 | 5.60 |
| *Dactylis glomerata* | DG | 80.80 | 68.00 | 68.80 | 61.60 |
| *Dianthus pontederae* | DP | 94.40 | 95.20 | 96.80 | 95.20 |
| *Echium vulgare* | EV | 23.20 | 18.40 | 25.60 | 21.60 |
| *Festuca rupicola* | FR | 64.95 | 49.87 | 52.40 | 61.81 |
| *Festuca vaginata* | FVg | 77.70 | 42.40 | 34.67 | 43.33 |
| *Festuca wagneri* | FW | 71.46 | 78.27 | 75.20 | 79.20 |
| *Filipendula vulgaris* | FiV | 16.00 | 24.80 | 4.80 | 5.60 |
| *Gypsophila paniculata* | GP | 80.00 | 70.40 | 56.00 | 44.00 |
| *Holoschoenus romanus* | HoR | 61.60 | 65.60 | 80.00 | 72.80 |
| *Hypericum perforatum* | HP | 40.80 | 50.40 | 42.40 | 47.20 |
| *Hypochoeris maculata* | HM | 86.40 | 84.80 | 80.00 | 72.00 |
| *Knautia arvensis* | KA | 15.20 | 8.00 | 4.80 | 3.20 |
| *Linaria vulgaris* | LVu | 21.60 | 18.40 | 18.40 | 19.20 |
| *Linum austriacum* | LA | 89.60 | 88.80 | 92.00 | 84.00 |
| *Lotus corniculatus* | LC | 62.40 | 52.80 | 35.20 | 43.20 |
| *Lythrum virgatum* | LyV | 67.20 | 74.40 | 72.80 | 62.40 |
| *Medicago lupulina* | ML | 93.60 | 89.60 | 84.00 | 87.20 |
| *Onobrychis arenaria* | OA | 80.00 | 75.20 | 68.80 | 80.00 |
| *Plantago lanceolata* | PL | 13.60 | 10.40 | 13.60 | 20.80 |
| *Poa angustifolia* | PoA | 94.40 | 89.60 | 90.40 | 87.20 |
| *Potentilla argentea* | PAr | 76.00 | 76.00 | 68.80 | 80.80 |
| *Potentilla recta* | PR | 44.80 | 50.40 | 44.00 | 43.20 |
| *Salvia pratensis* | SaP | 5.60 | 0.00 | 2.40 | 1.60 |
| *Scabiosa ochroleuca* | SO | 58.51 | 24.00 | 11.88 | 10.00 |
| *Senecio jacobaea* | SJ | 20.00 | 21.60 | 12.00 | 7.20 |
| *Silene conica* | SC | 88.80 | 89.60 | 90.40 | 89.60 |
| *Silene vulgaris* | SV | 85.60 | 85.60 | 71.20 | 58.40 |
| *Stachys germanica* | StG | 9.60 | 5.60 | 11.20 | 6.40 |
| *Stachys palustris* | StP | 29.60 | 16.80 | 0.80 | 0.00 |
| *Teucrium chamaedrys* | TeC | 10.40 | 12.80 | 13.60 | 8.80 |
| *Thymus glabrescens* | TG | 58.40 | 37.60 | 25.60 | 16.00 |
| *Tragopogon dubius* | TD | 75.80 | 72.80 | 24.00 | 13.60 |
| *Trifolium montanum* | TM | 45.60 | 32.80 | 34.58 | 32.00 |
| *Trifolium repens* | TR | 76.80 | 72.00 | 63.20 | 61.60 |
| *Verbascum phoeniceum* | VP | 99.20 | 91.20 | 97.60 | 96.00 |

Supplementary Table 5. Mean germination uncertainty (U) of the controls of the four germination cycles. The darkening hue of cells indicates the increasing mean U, different hues indicate significant differences (*p* ≤ 0.05). NA in the cells indicate the case when germination was not sufficient to calculate U.

| Species name | Abbreviation | K1 | K2 | K3 | K4 |
| --- | --- | --- | --- | --- | --- |
| *Achillea millefolium* | AM | 0.00 | 0.00 | 0.14 | 0.12 |
| *Agrimonia eupatoria* | AE | NA | NA | NA | NA |
| *Anthyllis vulneraria* | AV | 0.25 | 0.20 | 0.18 | 0.32 |
| *Astragalus cicer* | AC | 1.18 | 1.10 | 0.98 | 1.13 |
| *Betonica officinalis* | BO | 1.04 | 1.49 | 1.25 | 0.73 |
| *Brachypodium pinnatum* | BP | 1.28 | 0.95 | 1.35 | 1.03 |
| *Centaurea jacea* ssp. *jacea* | CJJ | 0.69 | 0.88 | 1.02 | 1.24 |
| *Centaurea jacea* ssp. *pannonica* | CJP | 0.55 | 0.96 | 0.74 | 0.77 |
| *Chrysopogon gryllus* | CG | 0.50 | 0.38 | NA | 0.18 |
| *Dactylis glomerata* | DG | 1.28 | 0.93 | 1.28 | 1.00 |
| *Dianthus pontederae* | DP | 0.29 | 0.10 | 0.10 | 0.24 |
| *Echium vulgare* | EV | 0.48 | 0.76 | 0.86 | 0.48 |
| *Festuca rupicola* | FR | 1.05 | 0.61 | 0.97 | 0.93 |
| *Festuca vaginata* | FVg | 1.17 | 1.17 | 0.60 | 0.78 |
| *Festuca wagneri* | FW | 0.96 | 0.73 | 1.28 | 0.73 |
| *Filipendula vulgaris* | FiV | 0.48 | 0.88 | 0.00 | 0.16 |
| *Gypsophila paniculata* | GP | 0.36 | 0.11 | 0.55 | 0.97 |
| *Holoschoenus romanus* | HoR | 1.31 | 1.17 | 1.09 | 1.19 |
| *Hypericum perforatum* | HP | 1.41 | 1.34 | 0.94 | 1.25 |
| *Hypochoeris maculata* | HM | 0.88 | 0.52 | 0.74 | 0.51 |
| *Knautia arvensis* | KA | 0.84 | 0.14 | 0.00 | NA |
| *Linaria vulgaris* | LVu | 0.89 | 0.38 | 0.13 | 0.19 |
| *Linum austriacum* | LA | 1.49 | 0.93 | 1.19 | 1.10 |
| *Lotus corniculatus* | LC | 0.76 | 0.16 | 0.40 | 0.77 |
| *Lythrum virgatum* | LyV | 1.19 | 1.36 | 1.21 | 1.32 |
| *Medicago lupulina* | ML | 0.10 | 0.00 | 0.00 | 0.16 |
| *Onobrychis arenaria* | OA | 0.56 | 0.18 | 0.55 | 0.49 |
| *Plantago lanceolata* | PL | 0.63 | 0.38 | 0.45 | 0.50 |
| *Poa angustifolia* | PoA | 1.08 | 1.06 | 0.54 | 0.67 |
| *Potentilla argentea* | PAr | 0.88 | 0.44 | 0.91 | 0.88 |
| *Potentilla recta* | PR | 0.76 | 0.67 | 0.29 | 0.44 |
| *Salvia pratensis* | SaP | NA | NA | NA | NA |
| *Scabiosa ochroleuca* | SO | 0.59 | 0.67 | 0.33 | 0.38 |
| *Senecio jacobaea* | SJ | 1.05 | 1.11 | 0.59 | 0.36 |
| *Silene conica* | SC | 0.23 | 0.18 | 0.72 | 0.52 |
| *Silene vulgaris* | SV | 0.81 | 0.62 | 1.28 | 1.20 |
| *Stachys germanica* | StG | 0.32 | 0.20 | 0.53 | 0.18 |
| *Stachys palustris* | StP | 1.36 | 1.35 | NA | NA |
| *Teucrium chamaedrys* | TeC | 0.55 | 0.68 | 0.98 | 0.40 |
| *Thymus glabrescens* | TG | 0.57 | 0.88 | 1.14 | 0.70 |
| *Tragopogon dubius* | TD | 1.45 | 1.32 | 0.99 | 0.87 |
| *Trifolium montanum* | TM | 0.26 | 0.18 | 0.47 | 0.18 |
| *Trifolium repens* | TR | 0.57 | 0.41 | 0.85 | 0.62 |
| *Verbascum phoeniceum* | VP | 0.26 | 0.30 | 0.84 | 0.31 |

Supplementary Figure 2. Differences in the Germination success (%, y axis of first and second column) and Germination uncertainty (U, y axis of third and fourth column) of the 44 studied species between the controls of the four germination cycles. Medians are displayed with solid black lines and means with dashed red lines. Where U could be not calculated boxes of the given treatment or even species are not shown (see Supplement Table 5). Lower-case letters indicate significant differences between the treatments (Tukey-test, p<0.05).

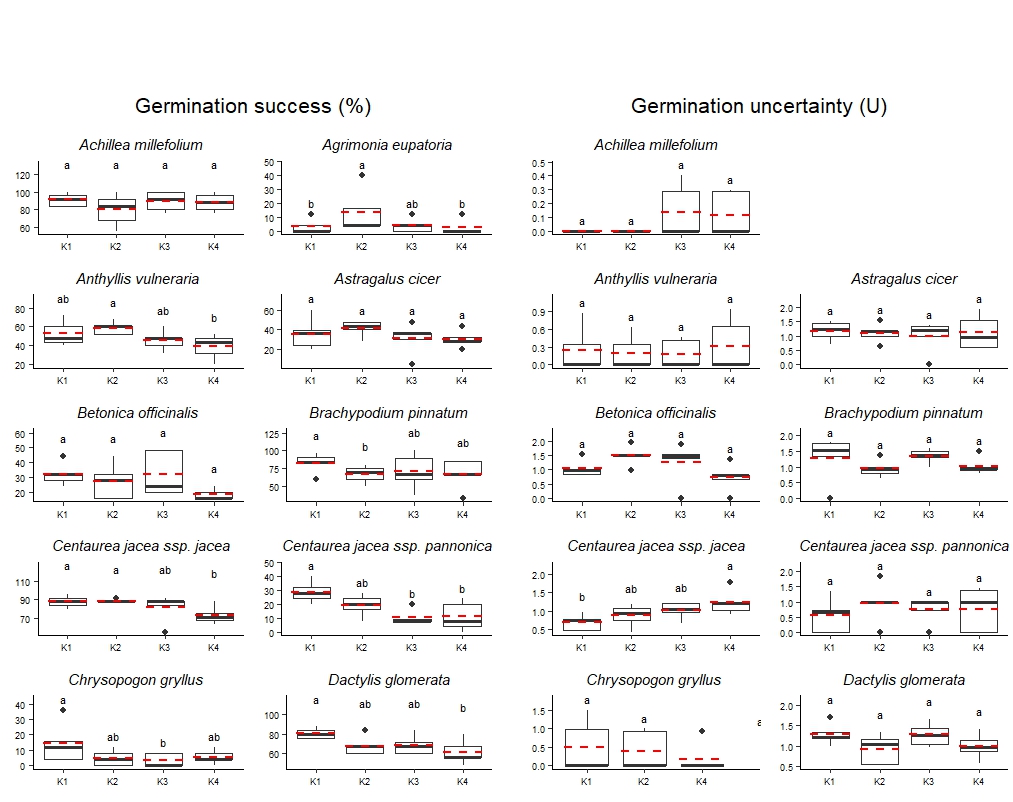

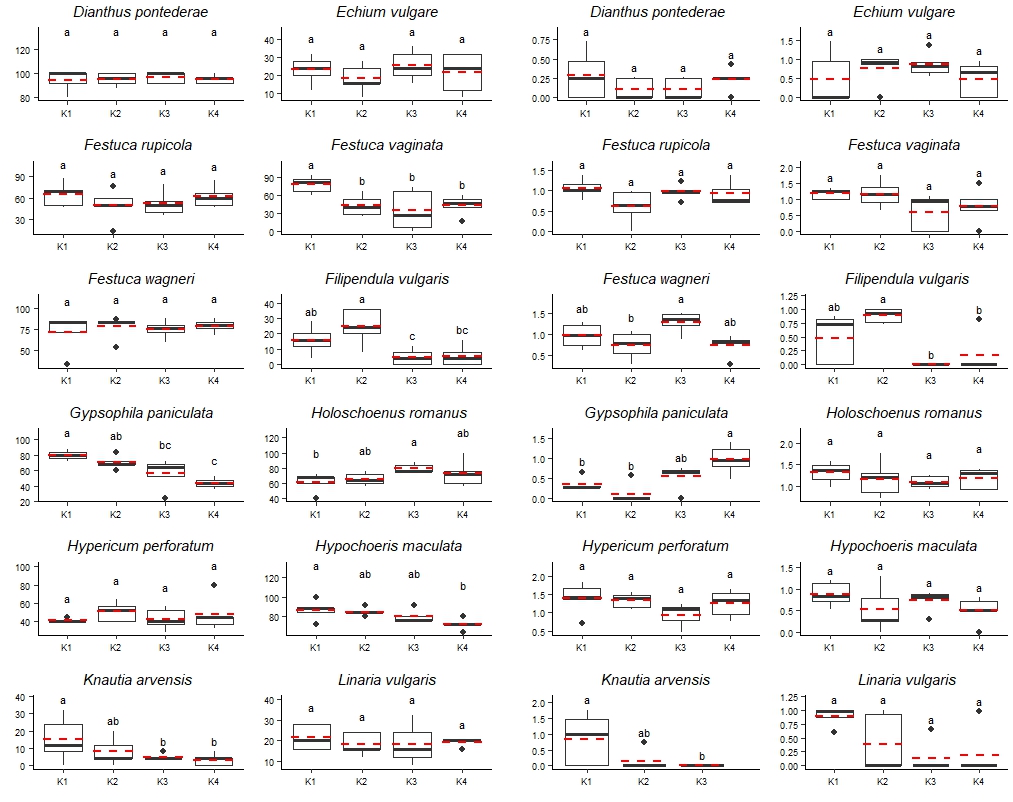

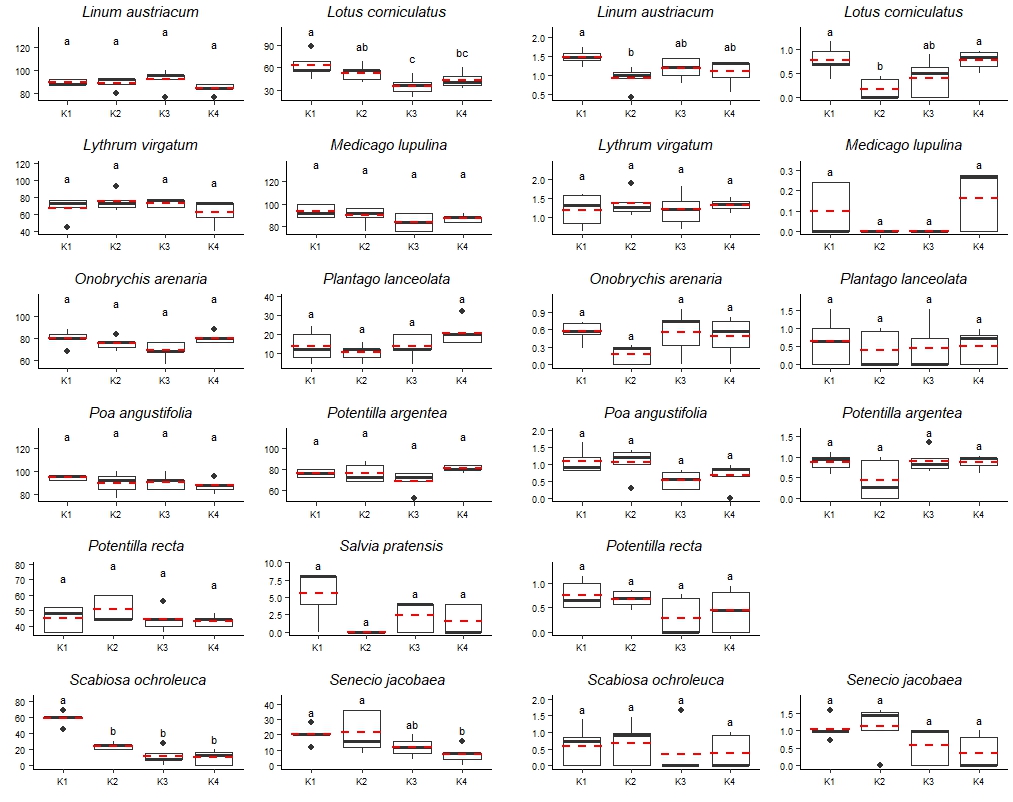

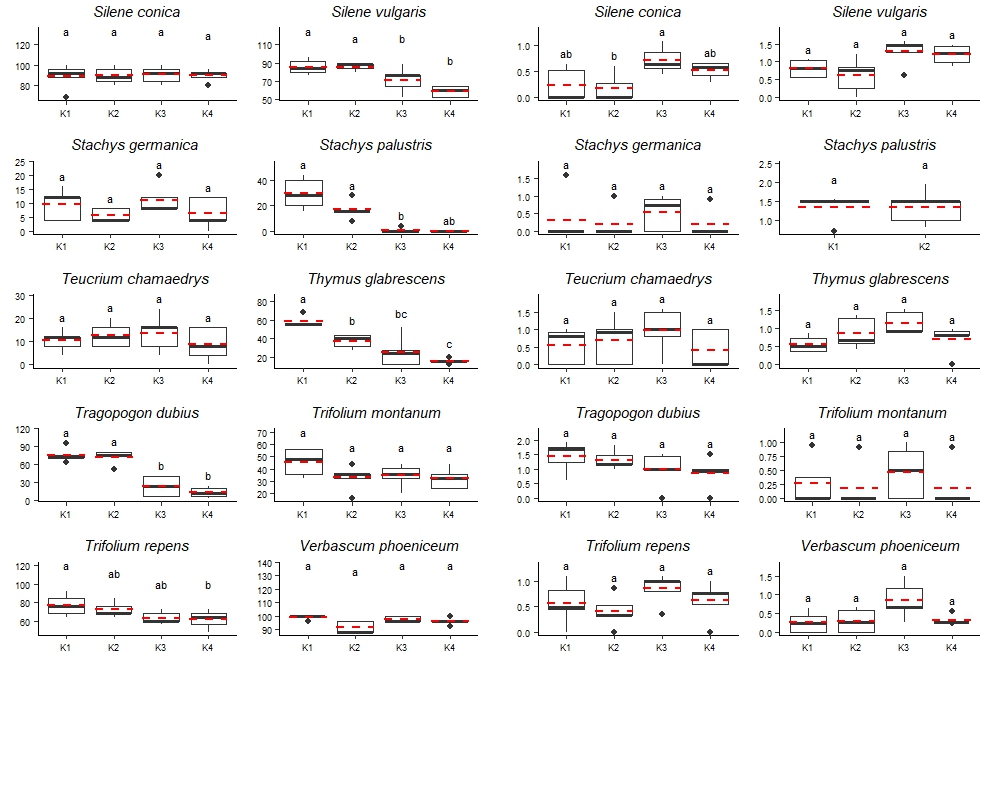
